# On the intrinsic curvature of animal whiskers

**DOI:** 10.1101/2022.05.18.492449

**Authors:** Yifu Luo, Mitra J.Z. Hartmann

## Abstract

Facial vibrissae (whiskers) are thin, tapered, and flexible hair-like structures that are an important source of tactile sensory information for many species of animals. In contrast to insect antennae, whiskers have no sensors along their lengths. Instead, when a whisker touches an object, the resulting deformation is transmitted to mechanoreceptors in a follicle at the whisker base. Previous work has shown that the mechanical signals transmitted along the whisker will depend strongly on the whisker’s geometric parameters, in particular on its taper (how diameter varies with arc length) and on its intrinsic curvature. Although previous studies have largely agreed on how to define taper, multiple approaches have been used to quantify intrinsic curvature. The present work, compares and contrasts different mathematical methods to quantify intrinsic curvature, including polynomial, fractional exponent, elliptical, and Cesàro. Comparisons are performed across ten species of whiskered animals, ranging from rodents to pinnipeds. The fractional exponent model is shown to be a particularly promising approach for distinguishing between whiskers of different species. We conclude with a discussion of the advantages and disadvantages of using the different models for different modeling situations.

## Introduction

For most mammals, whiskers (vibrissae) are an important source of tactile sensory information, enabling exploratory behaviors that range from climbing [1] to locomotion [2], to hunting [3]. Whiskers emerge from richly innervated follicles in the cheek and are often arranged in a regular array of rows and columns. Unlike insect antennae, whiskers have no sensory receptors along their length. Instead, when a whisker is deflected – either by direct physical contact or through fluid flow – the whisker transmits the mechanical deformation to its base [4-7]. The mechanical deformation is transduced into electric signals by different types of mechanoreceptors inside the follicle [8, 9].

The geometry and material properties of a whisker will determine its mechanical response and thus the information conveyed to the animal’s nervous system. Both whisker bending and vibration are essential for whisker function [5, 10-13], and these mechanical responses depend strongly on taper [5, 14-18] and intrinsic curvature [4, 19].

The taper of a whisker is well-defined in the literature: it describes how the whisker’s diameter changes along the arc length. As a first approximation, the diameter of a whisker tapers linearly from base to tip [15, 20-23]. Higher resolution measurements have refined this approximation: mouse whiskers were found to taper slightly more steeply in proximal regions (closer to the base) [24]. A similar high-resolution study has not yet been performed for rat whiskers.

In contrast to taper, which is well-defined, the intrinsic curvature of a whisker is more challenging to quantify. Approaches towards quantifying curvature have differed across studies and curvature lacks a unified representation. Previous studies have approximated the intrinsic curvature of a rat whisker in a variety of ways: as a quadratic Bézier curve [25], as a piecewise polynomial function [26], as a parabolic function (y=Ax^2^+Bx+C) [27], as a parabolic function with only a quadratic term (y=Ax^2^) [28, 29], and as a fifth-degree polynomial equation [14]. One recent study showed that the shape of rat whiskers can be mathematically transformed to fit intervals on the standard Euler spiral (κ=ŝ, where ŝ is the transformed arc length coordinate) [30], an approach which was later extended to include several other species [31].

The present work compares and contrasts different mathematical approaches towards describing whisker intrinsic curvature, and compares these approaches across ten different species: rat, mouse, gerbil, chinchilla, ground squirrel, rabbit, cat, harbor seal, sea lion, and pig. We discuss the advantages and disadvantages of the each of the approaches.

## Materials and Methods

Ethics statement: All experiments involving animals were approved in advance by the Institutional Animal Care and Use Committee of Northwestern University. All work on tissue from marine mammals was performed with authorization from the National Oceanic and Atmospheric Administration’s (NOAA’s) National Marine Fisheries Service (NMFS) in accordance with Marine Mammal Protection Act regulations.

### Data collection

A total of 1,762 whiskers from 10 species were used in this study (Table 1), including rats (*Rattus norvegicus*), mice (*Mus musculus*), gerbils (*Meriones unguiculatus*), chinchillas (*Chinchilla chinchilla*), ground squirrels (*Ictidomys tridecemlineatus*), rabbits (*Oryctolagus cuniculus*), cats (*Felis catus*), harbor seals (*Phoca vitulina*), sea lions (*Zalophus californianus*), and pigs (*Sus domesticus*).

**Table 1.** The number of whiskers and specimens used in this study.

| Species | # of individuals | # of whiskers | Sex and age |
| --- | --- | --- | --- |
| Rat | 4 | 225 | All female, 5~36 months old |
| Mouse | 8 | 323 | All male, 6~8 weeks |
| Gerbil | 3 | 172 | 1 male, 2 female, 3 months old |
| Chinchilla | 2 | 95 | Unknown |
| Ground squirrel | 3 | 116 | Mixed sex, adult |
| Rabbit | 2 | 126 | Female, 10 months |
| Cat | 5 | 228 | Female, 11 months |
| Harbor seal | 6 | 306 | 2 male, 4 female, all >1yr old |
| Sea lion | 2 | 148 | Female, subadult and 5+ years old |
| Pig | 1 | 23 | Female, age unknown |

Whiskers from harbor seals were obtained from animals post-mortem in collaboration with Allied Whale at the College of the Atlantic (Bar Harbor, ME). For all other species, whiskers were taken from freshly euthanized laboratory animals that had been used in unrelated experiments. Each whisker was identified by its row and column identity, carefully plucked with fine forceps at its base, and then scanned in 2D on a flatbed scanner (Epson Perfection 4180 Photo) at 6,400 dpi resolution. Whisker shape was extracted as a 2D curve from the scanned image using custom-written software in MATLAB.

To reduce pixelation noise, each extracted whisker curve was smoothed using local regression (MATLAB “loess”) on a moving window of size 1,000 pixels (∼4 mm). The whisker curve was then resampled into equally spaced nodes (0.25 mm) for final analysis. In previous studies of rat whisker curvature [27-29], only the proximal 60-70% of the whisker was considered planar and used to quantify curvature. In contrast, the present work used the total length whiskers for all species in all analysis.

### Defining “standard orientation” for Cartesian descriptions of whisker shape and the whisker arc length

Although some representations of whisker curvature are coordinate free, others rely on a choice about how to place the whisker in a Cartesian coordinate system. We defined the “standard orientation” of a 2D whisker in Cartesian coordinates as follows: the whisker base (the first node) was placed at the origin and the proximal 15% of the whisker was aligned with the positive x-axis. The whisker was oriented such that most whisker nodes are in the first quadrant; in practice this meant that whiskers were oriented “concave upwards.” All whiskers were placed in standard orientation before curve fitting.

The arc length of a whisker was defined as the summation of the lengths of the segments formed by adjacent nodes along the trace, as distinct from the straight-line base-to-tip length.

Fig 1 shows all whiskers in standard orientation, grouped by species. The inset histograms show statistics of whisker length.

**Fig 1.**
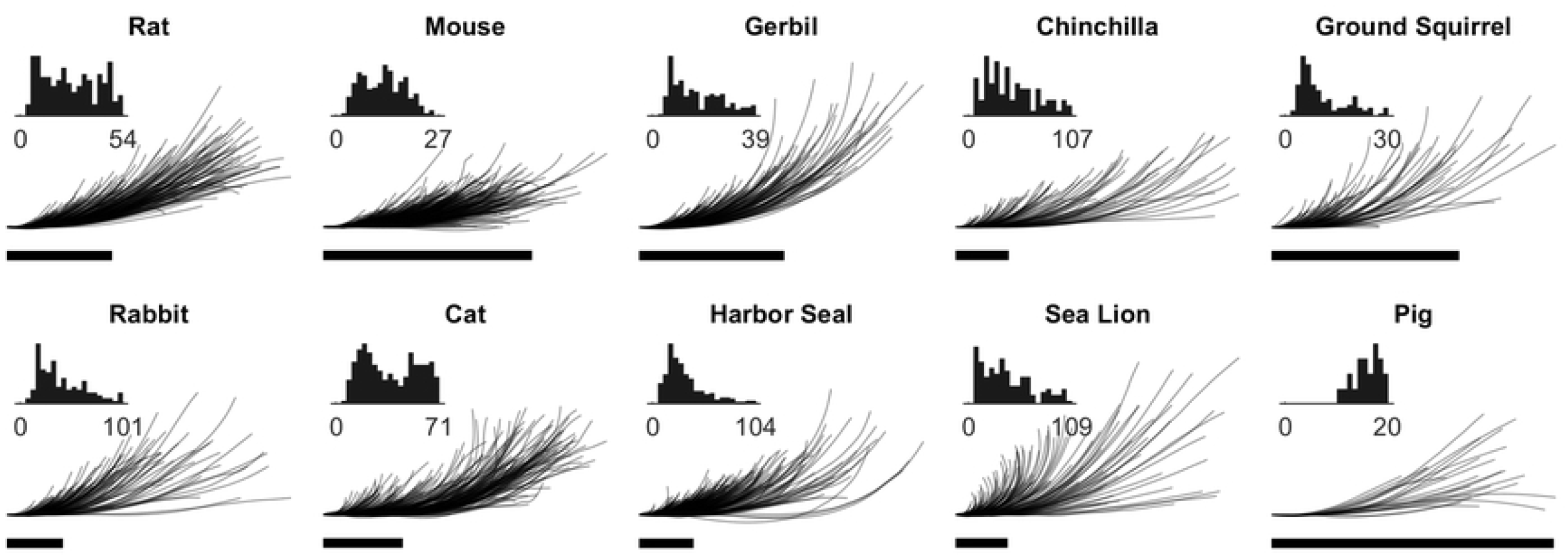
All whiskers are placed in standard orientation before curve fitting. The proximal 15% of each whisker is aligned with the positive x-axis (horizontal). Scale bar: 20 mm. Inset: histogram of whisker length (unit: mm).

### Total least squares

Total least squares was used as a metric to account for observational error in both x and y dimensions of the variables. The residual sum of squares (RSS) was used as an optimality criterion in parameter selection and model selection. For a set of n observations (x_i_, y_i_), i ≤ n, RSS is calculated as:

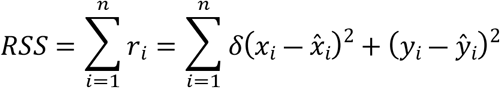

where 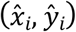 is the predicted value of observation (x_i_, y_i_), and δ=Var(y_i_)/Var(x_i_). (Var(·): variation)

In practice, the residual r_i_ (that is, the distance between the observed data and the fitted curve) is approximated by 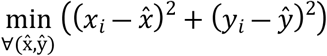. The total least squares metric was used for all models (both linear and non-linear) unless otherwise specified.

## Results

In the following sections, we compare and contrast multiple approaches towards quantifying whisker curvature.

### Polynomial model

Polynomial equations were fit to each whisker from base to tip. This approach relies on orienting the whisker in a Cartesian coordinate system. Because each whisker was placed in standard orientation before curve fitting (see *Materials and Methods*), both the constant term a_0_ and the linear coefficient a_1_ are 0 by definition. Therefore, each whisker is fit with the following polynomial equation:

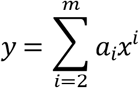

where m is the highest chosen order.

We noticed that when placed in standard orientation, the tips of some whiskers curved so much that they went backwards along the x-axis. In these cases, the whiskers were simply trimmed at the point where they began to curve backwards. Notice that this cut-off is a disadvantage of the polynomial model, one that is avoided by the more complex models described in subsequent sections.

To prevent overfitting, we tested a variety of polynomial models of order increasing from 2. All whiskers were fit by the model y=a_2_x^2^, y=a_3_x^3^+a_2_x^2^, y=a_4_x^4^+a_3_x^3^+a_2_x^2^, and y= a_5_x^5^+a_4_x^4^+a_3_x^3^+a_2_x^2^, separately. For each model, the model coefficients and the RSS were obtained for each whisker.

Fig 2 shows, for all species, how the RSS for each whisker changes with increased order of polynomial models. To avoid overfitting, a higher order polynomial model is rejected when it does not eliminate 90% or more of the variation of the previous model. Summarizing the results of Fig 2, a quadratic model y=a_2_x^2^ is sufficient for rats, mice, and pigs, but all other species require y=a_3_x^3^+a_2_x^2^. Any polynomial model with order higher than 3 is an overfitted model.

**Fig 2.**
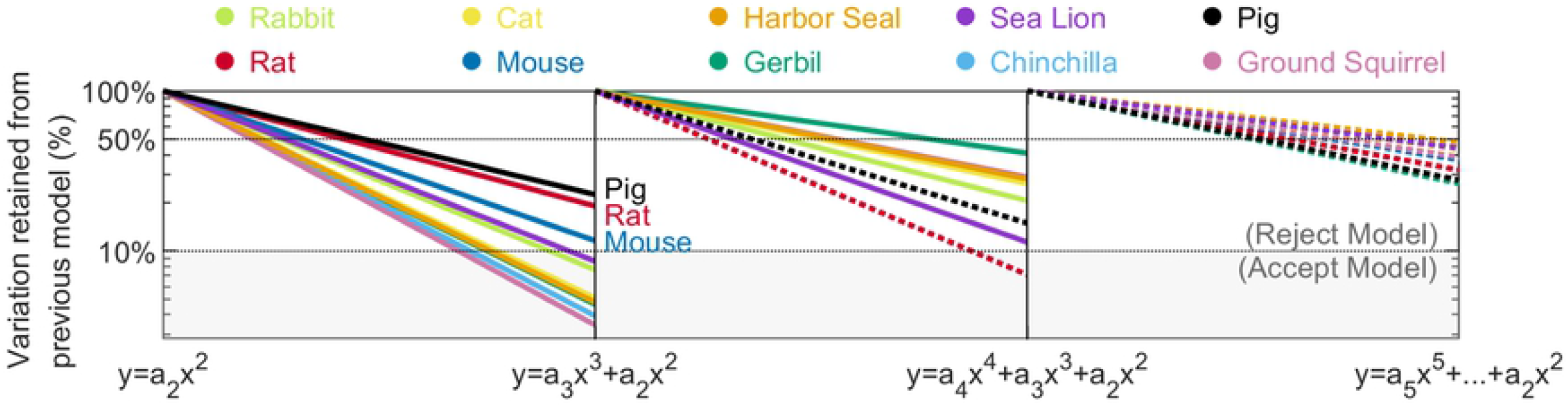
Decreases in the RSS from each lower order polynomial model to each higher order polynomial model are shown as a percentage of the lower order model (normalized to 100%). If a higher order model retains less than 10% of variation from the lower order model, it is accepted, otherwise it is rejected. Within each panel of the figure, species with solid lines are evaluated to determine if they would be overfit by the subsequent model. In the first panel, all lines are solid because all species are evaluated. The species pig, rat, and mouse are determined to be well fit by the model y= a_2_x^2^ and overfit by the model y=a_3_x^3^+a_2_x^2^. These three species are shown as dotted lines in the subsequent panel, while the remaining seven species are shown as solid lines. These seven species are well fit by the model y=a_3_x^3^+a_2_x^2^ but overfit by y=a_4_x^4^+a_3_x^3^+a_2_x^2^ and therefore shown as dotted lines in the third panel.

### Fractional exponential model

It is notable that the polynomial description of whisker curvature requires both quadratic and cubic terms for most species. Upon closer examination, it can be shown that these two terms contribute in different ways to whisker fitting. In Fig 3AB, the residuals of the whisker fit are plotted as histograms for the models y=a_2_x^2^, y=a_3_x^3^, and y=a_3_x^3^+a_2_x^2^. The figure shows clearly that the residual histogram is biased in opposite directions for y=a_2_x^2^ and y=a_3_x^3^, and that the bias is much smaller for the polynomial model y=a_3_x^3^+a_2_x^2^, which combines the two terms.

**Fig 3.**
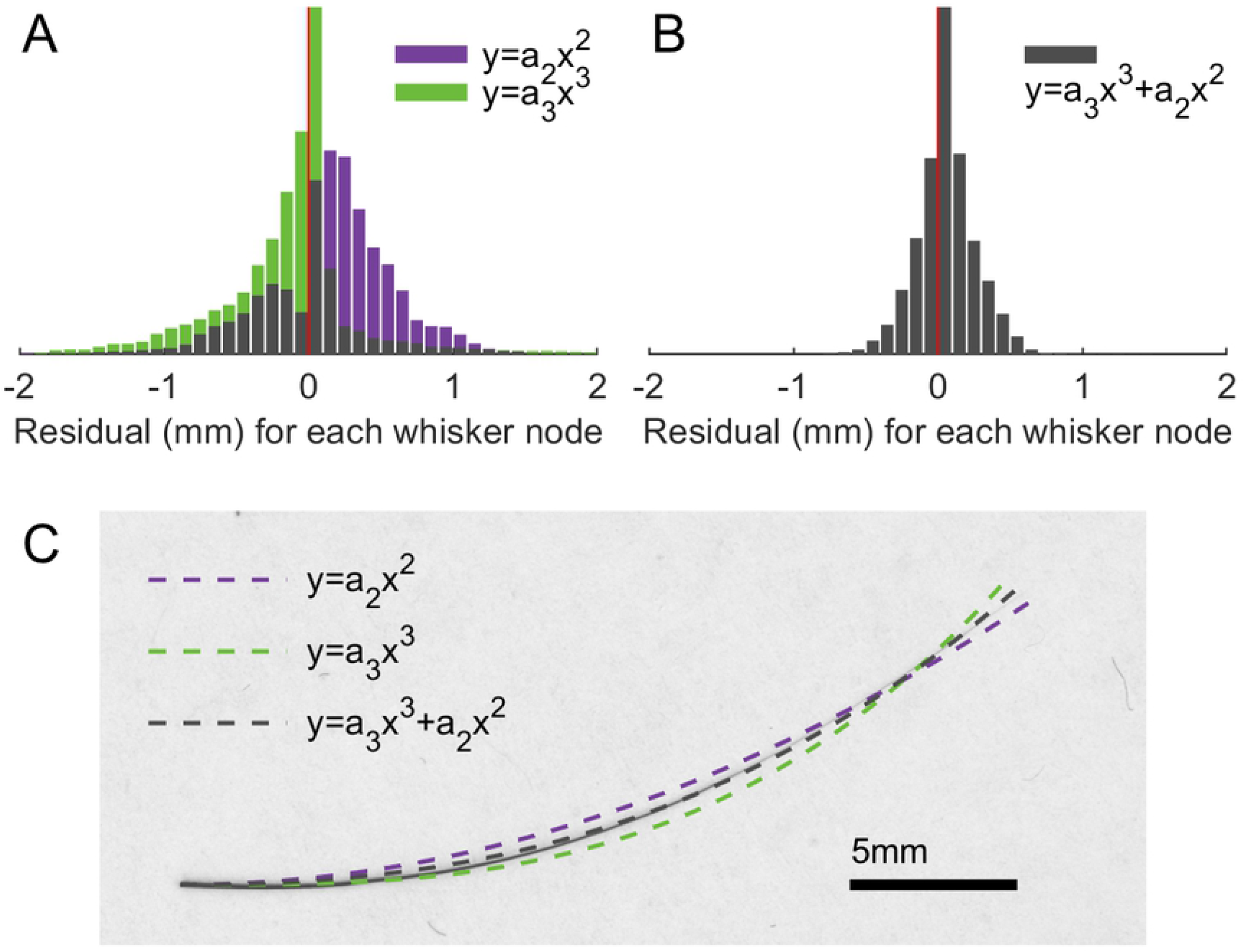
The quadratic and cubic terms in a polynomial model make different contributions to the fit. **(A)** The histograms of residuals for rat whiskers using the model y=a_2_x^2^ (purple) and y=a_3_x^3^ (green) are skewed to the opposite sides of the middle line at 0. **(B)** The histogram of residuals for rat whiskers using the model y=a_3_x^3^+a_2_x^2^ is shown; skewness is near zero. **(C)** Different curves fits are plotted for an exemplar scanned rat whisker. It can be observed that the quadratic and cubic models generate residuals on opposite sides of the whisker. The whisker shape lies between the two well-fit single-term curves, y=a_2_x^2^ and y=a_3_x^3^.

Geometric intuition for these biased residuals can be obtained by examining an exemplar whisker (Fig 3D). This whisker is placed in standard orientation, and its shape falls between the two best-fit single-term polynomial curves, y=a_2_x^2^ and y=a_3_x^3^. A polynomial model combining the two terms provides a better fit – in other words, whisker shape is neither quadratic nor cubic.

Based on these results, we hypothesized that a model y=a_β_x^β^ with a fractional exponent β between 2 and 3 would better describe the general shape of a whisker than a polynomial model limited to integer exponent values. We hypothesized further, that the parameter β might provide a useful basis to distinguish between whiskers from different species. This analysis was performed through the following steps.

#### Step 1: Optimize fractional exponent values for each whisker

To find the model coefficient a_β_ and the fractional exponent β value for each whisker, we ran a nested optimization. In the inner optimization, the coefficient a_β_ was optimized based on the least RSS 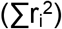 given a fixed value for β. In the outer optimization, the optimal β value was chosen based on a slightly different parameter, namely, the overall minimal square of residual sum (Σr_i_)^2^. This metric accounts for bias by introducing product terms.

In Fig 4A, the optimized fractional exponent β for each whisker is shown in a heatmap, with each row showing results for each species. The fractional exponent values are mostly distributed between 2 and 3, confirming our hypothesis that the shape of most whiskers lies between quadratic and cubic curves and should be well approximated by the model y=a_β_x^β^ (2<β<3). However, Fig 4A also shows a non-negligible number of whiskers for each species with optimal values for β that are less than 2 or greater than 3. Whiskers with β values less than 2 are usually short and straight. Whiskers with β values greater than 3 have increasing rates of change of curvature especially approaching the tip.

**Fig 4.**
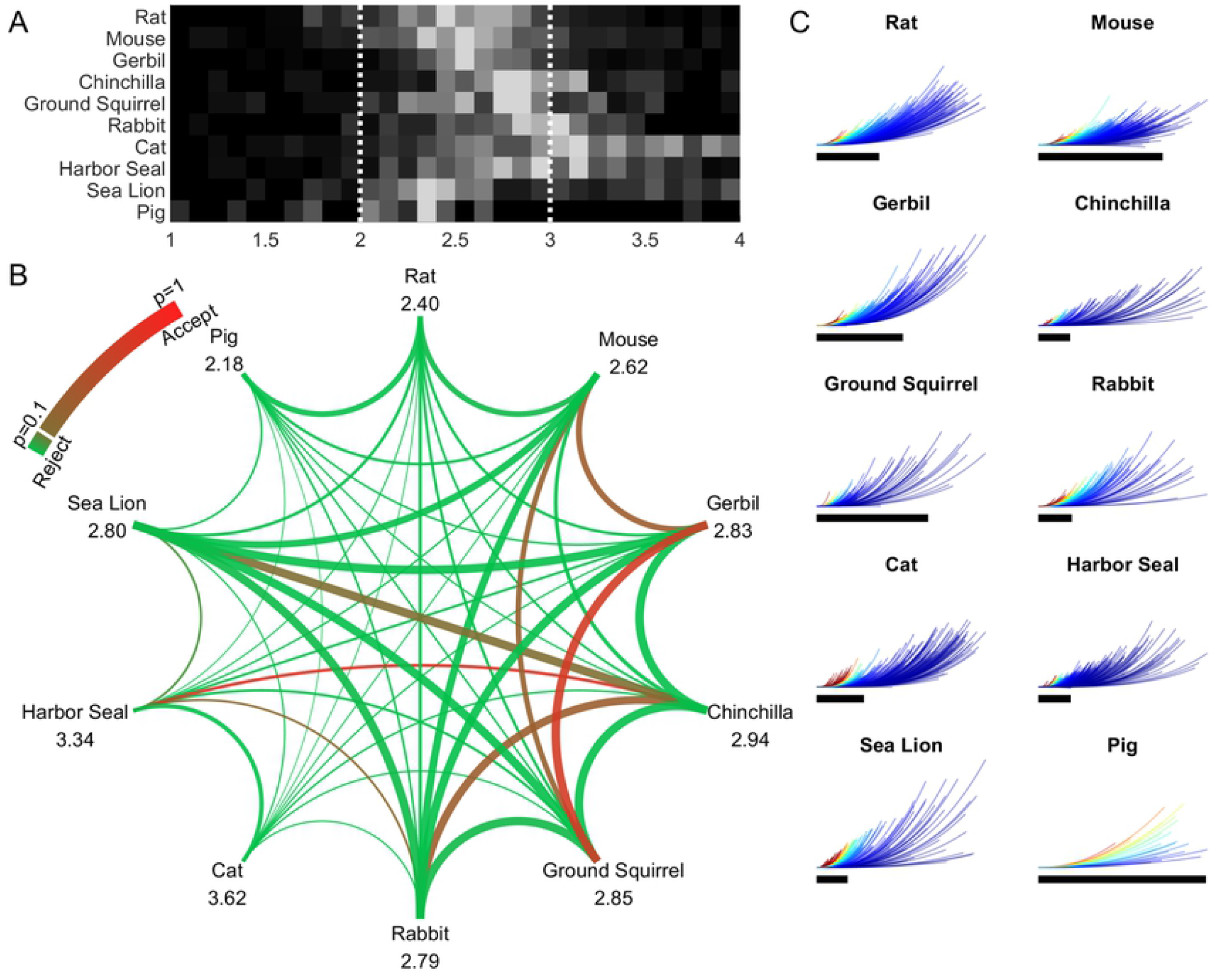
A fractional exponent model can be used to quantify whisker curvature. **(A)** A heatmap of exponent value for whiskers of all species is shown. **(B)** This graph shows results of a cross-species comparison generated from the unpaired t-tests in step 2. The number under each species name is the generalized fractional exponent β value determined in step 3. The color of the curve indicates the p-value of the student’s unpaired t-test. The green color rejects the null hypothesis, indicating that whiskers can be distinguished, while the red color accepts the null hypothesis, indicating that whiskers cannot be distinguished. The thickness of each line indicates the similarity of the two fractional exponent value optimized in step 3. **(C)** Whiskers are colored according to the magnitude of their a_β_ coefficients, using the generalized fractional exponent value.

#### Step 2: Quantify significance of cross-species differences

In this analysis, whiskers with values of β outside the 95% confidence interval for that species were excluded to reduce variability. The number of whiskers removed for each species is generally less than 5% of the total number. The significance of cross species differences was then determined by conducting a student’s unpaired t-test (α=0.05) between each pair of species. The null hypothesis is that the fractional exponent values of whiskers are from independent random samples from normal distributions with equal means and variances, that is, the whisker geometry is not separable between the two species.

Results of the hypothesis testing for all pairs are shown in Fig 4B. A total of 45 species pairs were tested. Using the value of the fractional exponent β as the sole parameter, 39 species pairs are separable (p<0.1), of which 36 pairs are strongly separable (p<0.01). Six species pairs are not separable: mouse vs. gerbil (p=0.30), mouse vs. ground squirrel (p=0.25), gerbil vs. ground squirrel (p=0.67), chinchilla vs. rabbit (p=0.29), chinchilla vs. harbor seal (p=0.78), rabbit vs. harbor seal (p=0.17).

#### Step 3: Optimize the value of the fractional exponent β across each species, so that *all* whiskers from the same species share the same β value

This analysis was again performed using a nested optimization. It first minimized the overall RSS 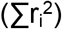 of all whiskers for a given S value. Then the optimal β value with the least overall square of residual sum (Σr_i_)^2^ was selected.

Note that in contrast to step 1, the fractional exponent model used in this step does not optimize the fit for each individual whisker, instead, it is optimized to describe the curvature of all whiskers of a species. Species that were found to be indistinguishable in step 2 also have similar generalized fractional exponent values. These values are reported at the nodes of the graph of Fig 4B.

Finally, having generalized the β coefficient for each species, we examined how the a_β_ coefficient varied within a single species. Fig 4C shows the distribution of the a_β_ coefficient among whiskers of the same species. The values of a_β_ tend to be different for short and long whiskers. Short whiskers tend to have larger and more variable values of a_β_ (as seen in the multiple colors identifying short whiskers), while longer whiskers have lower a_β_ values. These results are consistent with previous work [29] in which rat whiskers were fit to quadratic equations: a small change in the tip location for shorter whiskers will result in a large change in the a_β_ coefficient.

### Elliptical model

As indicated earlier, polynomial fits cannot accurately model whiskers whose tips go “backwards” (in the direction of the negative x-axis) when placed in standard orientation. We refer to these whiskers as “backwards whiskers.” Modeling the whisker as a segment of an ellipse can fit whiskers of this backwards type. We therefore considered the possibility that whisker curvature changes non-linearly from base to tip in a way that can be described by an elliptical model κ(x, y) = A^4^B^4^/(A^4^y^2^+B^4^x^2^)^3/2^. A point (x, y) is taken from a curve segment on an ellipse. The curve segment starts at the origin, initially moving along the positive x-axis, and going into the first quadrant. The equation for the ellipse is x^2^/A^2^+(y-B)^2^/B^2^=1.

This model has two parameters, A and B, corresponding to the lengths of major and minor axes, respectively. The model coefficients were fit individually for each whisker.

Similar to previous analyses, an optimization process was run to determine the two coefficients, minimizing the RSS between each whisker and the fit elliptical arc. Fig 5A shows some exemplar whiskers and their best-fit ellipses. Most whiskers are fit within a single quarter of the elliptical arc, and some backwards whiskers can be accurately fit by extending the arc beyond one quarter. Fig 5B shows histograms of the percentage of the quarter-ellipse that is being used to fit the whiskers for each species. Backwards whiskers are observed as those occupying more than 100% of the quarter-ellipse (to the right of the dotted vertical line). It is notable that the minimum of all histogram bin values is ∼40%, indicating that there are no cases in which a whisker was fit with a disproportionately large ellipse.

**Fig 5.**
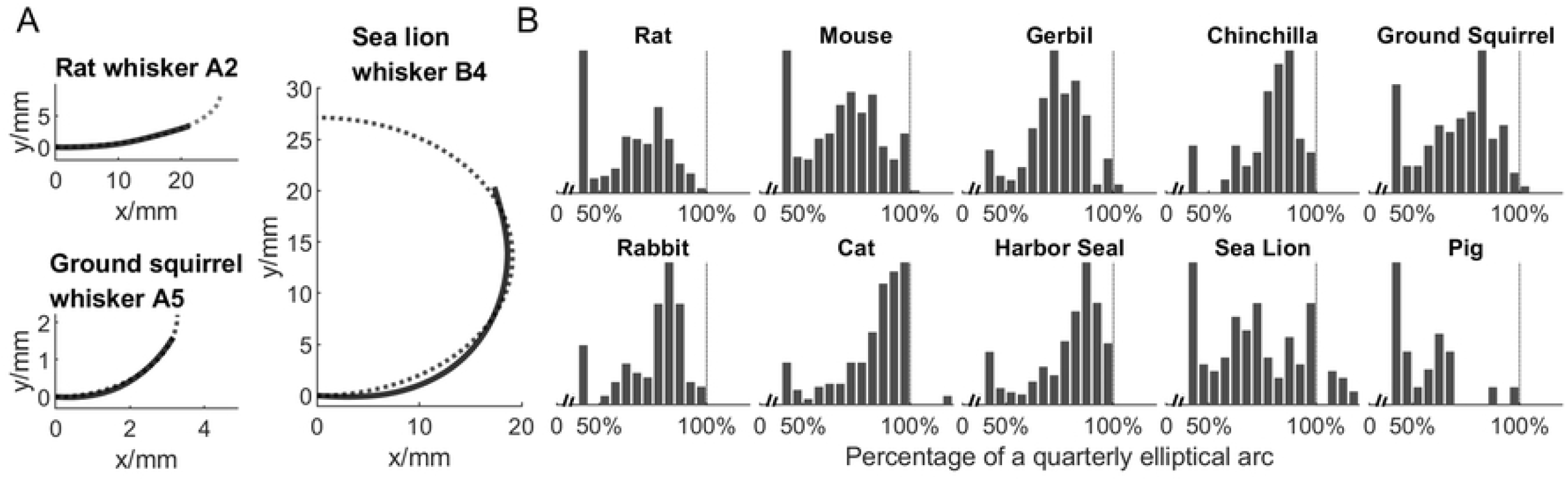
Results of fitting whiskers to an elliptical model are shown. **(A)** Three exemplar whiskers fit to ellipses are shown. Solid curve: whisker. Dotted curve: fit elliptical arc. **(B)** For each species, the histogram shows the number of whiskers that occupy a given percentage of a ¼ elliptical arc. Values greater than 100% (vertical dotted lines) indicate that the whisker occupied more than ¼ of the ellipse arc. The histogram bin value adds up to the total number of whiskers shown in Table 1.

### Linear curvature model: Cesàro equation

Both polynomial and elliptical models rely on the choice of whisker orientation in a Cartesian coordinate system. Several previous studies have described rat whisker shape using a coordinate-invariant representation, assuming that curvature (κ) changes linearly with arc length (s) [28, 30, 31]. In other words, these studies have assumed that κ(s)=As+B, where A and B are coefficients that can be different for each whisker. The sign of the coefficient A determines whether the curvature increases or decreases from a starting value B at the whisker base. We fit this model to the whiskers of all species in our dataset, optimizing the coefficients A and B by minimizing the summed Euclidian distance between each whisker and a curve of equal arc length generated by the equation κ(s)=As+B.

Whisker shapes are generally well represented by curves with a linear rate of change of curvature. Fig 6A shows two exemplar whiskers, from a cat and a sea lion. Linearly changing curvature can capture a wide variety of whisker shapes. With only two parameters, curve fitting works for whiskers that are approximately quadratic or cubic, as well as for whiskers that are “flat” at the beginning and curved up towards the tip (typical of many cat whiskers), and also for whiskers in which the tip goes backwards (typical of many sea lion whiskers).

**Fig 6.**
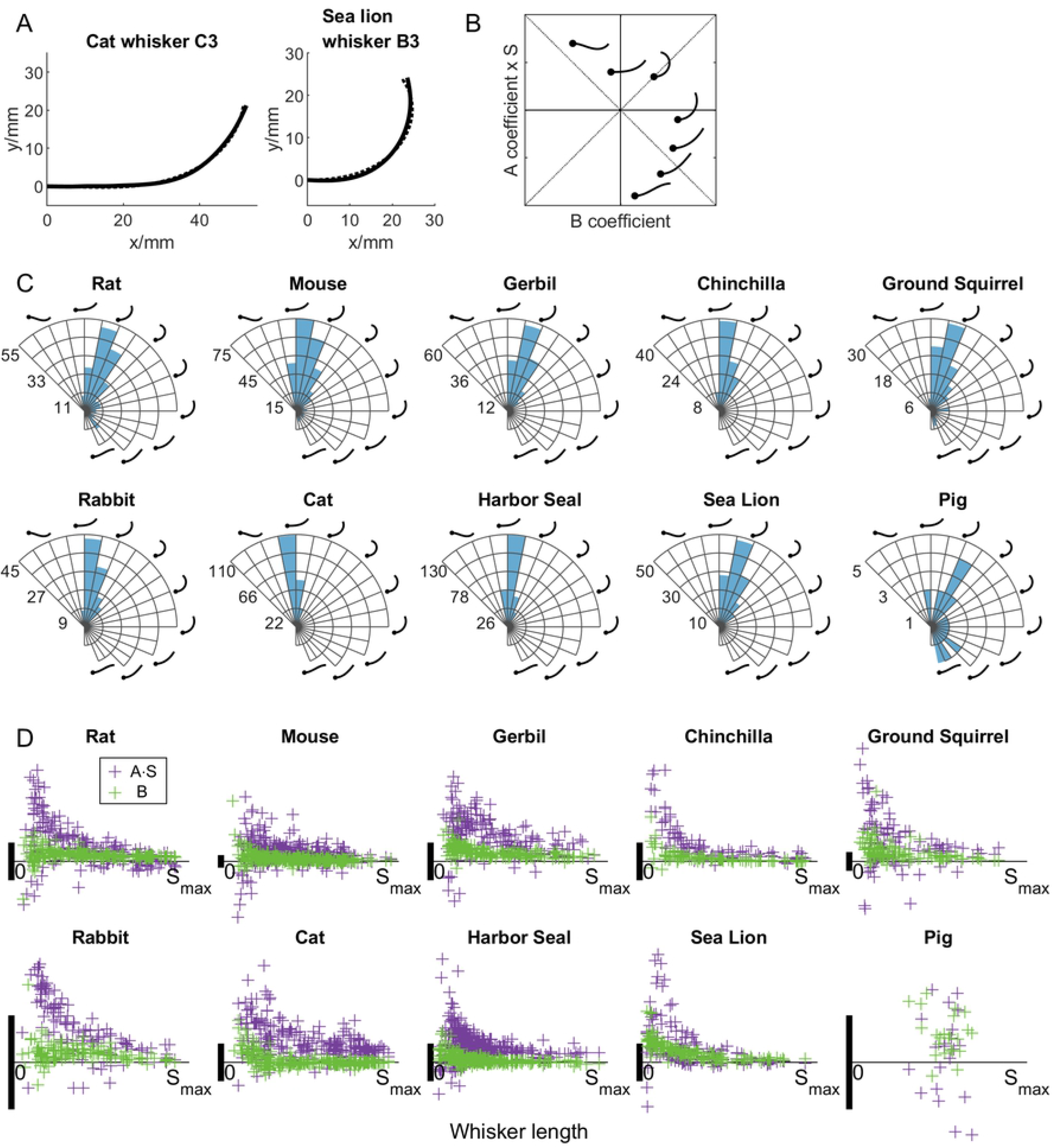
Whiskers can be fit to Cesàro curves. **(A)** Two exemplar whiskers fit to Cesàro equations are shown for cat and sea lion. Solid curve: whisker. Dotted curve: fit curve. **(B)** In the present dataset, whiskers could be classified into 9 different shapes, 7 of which are illustrated in the (B, A·S) coefficient space. **(C)** Polar histograms indicate the total number of whiskers in each region of the coefficient space for each species. The coordinates in each panel are the same as in (B). The radial axis indicates the number of whiskers. **(D)** The two coefficients B and A·S show large variance for short whiskers compared to long whiskers. In all plots the vertical scale bar represents 0.05mm.

Given the definition κ(s) =As+B, the general shape of a curve can be categorized into 16 types. In our dataset, we found that the shape of a whisker (from any species) was limited to nine of these 16. The relative magnitudes of the two coefficients determine the shape of the curve; seven distinct examples are schematized on the plot of coefficient space in Fig 6B.

After optimizing fits for all whiskers across all species, the majority (74.3%) of whiskers were found to fall into the first quadrant of the coefficient space, as shown in Fig 6C. For these whiskers, the curvature starts with a positive value at the whisker base (B>0) and increases towards the tip (A>0). A smaller number of whiskers (14.6%) were found to lie in the second quadrant (B<0, A>0), and fewer (11.0%) in the fourth quadrant (B>0, A<0), in which curvature gradually decreases from the base. In our dataset, only 2 out of 1762 whiskers were found to lie in the third quadrant. Across whiskers of all species, 80.3% have positive curvature all the way from whisker base to the tip.

We next examined whether the distribution of A and B coefficients were different for long and short whiskers. Fig 6D shows the coefficients plotted against whisker arc length. Across all species except the pig, shorter whiskers tend to have higher variability in both coefficients than longer whiskers. The large variation shown for short whiskers is also consistent with that shown for the fractional exponent model.

### Conforming whiskers onto a universal Euler spiral

Curves described by Cesàro equations (κ=As+B) are given the name “Euler spirals.” The particular curve defined by κ(s)=s (A=1, B=0) is referred to as the “universal Euler spiral.” A recent study [30] has shown that the shape of rat whiskers can be mathematically transformed onto arc segments of different lengths at different locations on the universal Euler spiral.

Fig 7A is replicates the results for rat whiskers shown in previous work ([30]). The mean locations of the conformed whiskers on the Euler spiral do not show strong correlations with either row (Pearson correlation test, ρ=-0.109) or column (ρ=0.102), or with whisker arc length (ρ=-0.141), or with the curvature averaged over the arc length (=A*S/2+B, where S is the total length of the whisker. ρ=0.152) (Fig 7BCDE). All of these quantities are mapped in a non-systematic way onto the universal Euler spiral. In other words, transforming the whiskers to lie on the Euler spiral loses information about whisker identity, whisker arc length, and average curvature over the arc length.

**Fig 7.**
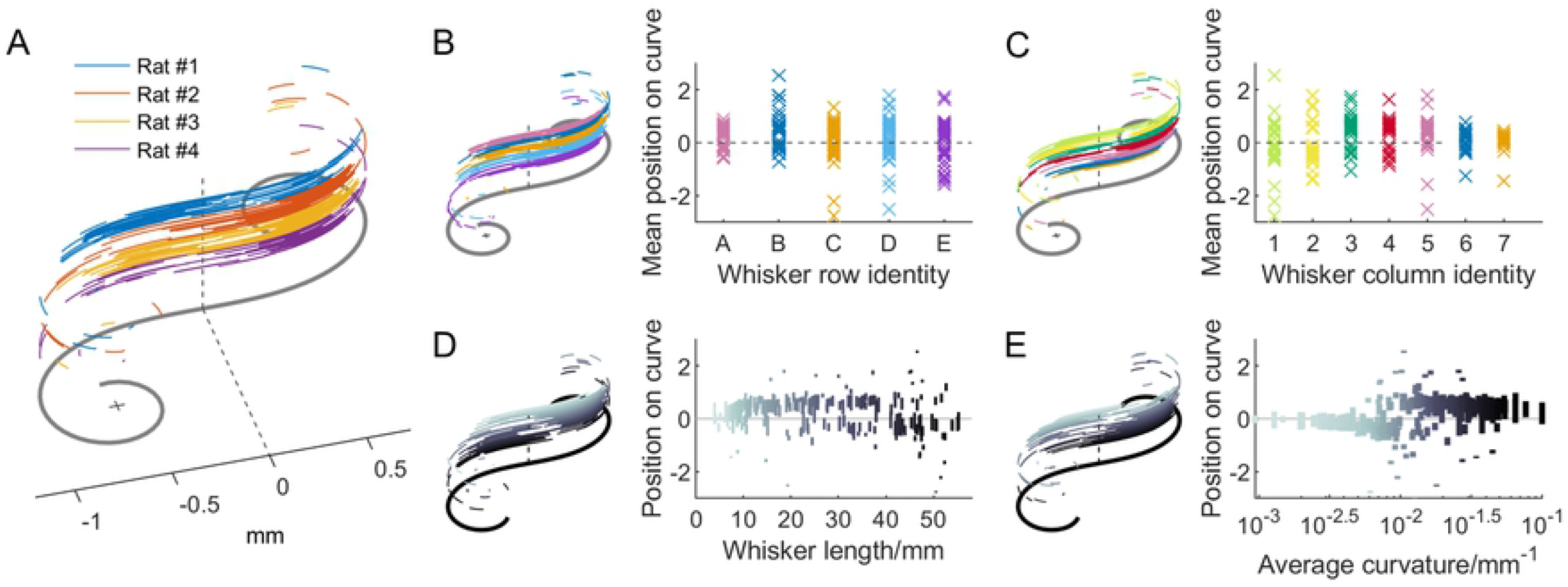
Whiskers are conformed onto the standard Euler spiral. (A) Individually layered 225 whiskers from 4 rats were conformed onto a standard Euler spiral shown underneath in gray. Whisker from each rat individual was colored distinctly. (B) Left: the whiskers are group by the row identity. Right: each data point shows the mean position of the conformed whisker on the standard Euler spiral. (C) The whiskers are group by the column identity. (D) Left: the whiskers are sorted by their arc length. Right: each vertical line shows the span of the conformed whisker on the standard Euler spiral. (E) The whiskers are sorted by their averaged curvature, defined as A*S/2+B.

Thus while it is true that each whisker can be mathematically transformed so as to lie on a universal Euler spiral, that fact would be true for any curve that follows κ=As+B (subtract B and divide by A), and the value of doing so is not immediately clear.

### Comparing modeling approaches

Fig 8 evaluates the different model types for all species using the RSS as a metric after removing outliers for each species (fewer than 3% on average). Unsurprisingly, models with two parameters tend to have lower values of the RSS than single parameter models. More specifically, the two-parameter polynomial model and the Cesàro fit always have a lower RSS than the best single parameter model. However, for 7/10 species, the elliptical fit model generates RSS values that are larger than the best single parameter model.

**Fig 8.**
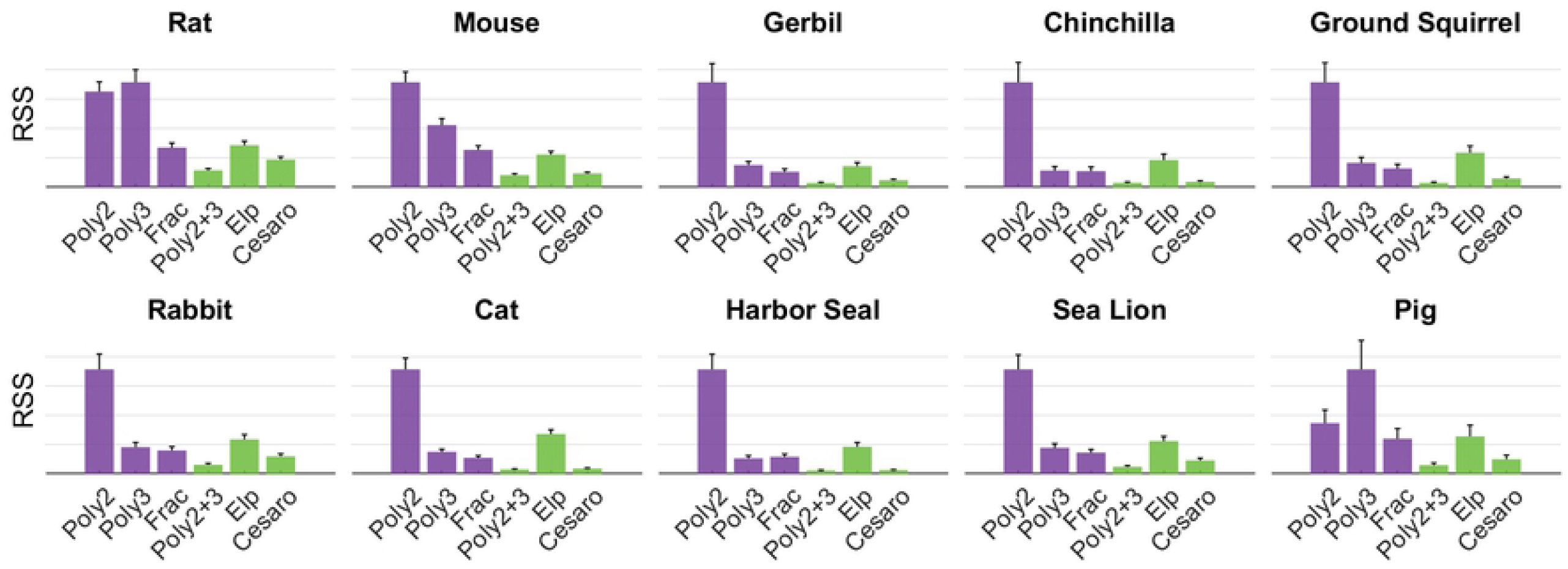
RSS is compared for all models for all species. Y-axis is normalized to the maximum value. All plots show adjusted RSS to account for differences in the number of parameters (either one or two). Poly2: y=a_2_x^2^. Poly3: y=a_3_x^3^. Poly2+3: y= a_3_x^3^+a_2_x^2^. Frac: fractional exponent model. Elp: elliptical model.

Within two parameter models, the polynomial model has lower RSS values than the elliptical model for all species, and a lower RSS value than the Cesàro model for all species except the harbor seal. Within single parameter models, the fractional exponent model has lower RSS values than the quadratic model for all species, and lower RSS values than the cubic model for all species except the sea lion.

Overall, the best single parameter model is the fractional exponent, while the best two-parameter model is either polynomial or Cesàro. The elliptical model does not appear to be a good choice for any species.

## Discussion

The present work has compared and contrasted different approaches for quantifying whisker curvature. Different approaches may be appropriate for answering different questions.

The advantage of a single-parameter polynomial model is that it is simple and extremely intuitive: it is easy to visualize a quadratic or a cubic curve. Single-parameter polynomial models generate reasonably good fits for all whiskers (Fig 2 and 8). However, a significant disadvantage is that it is not clear how to compare these fits across species.

The fractional exponent model fits whiskers with very low residuals, and in addition, a single parameter can be defined in a consistent way across whiskers across species. The fractional exponent approach is the only one that directly yields a single hyperparameter that can be compared across species. A second parameter then identifies each whisker.

The elliptical model (two parameters) does not appear to be a good choice to describe whisker shape. Although it can fit “backwards” whiskers, the residual values are high.

The Cesàro equation (also two parameters) generally yielded the best fits for all whiskers, and can also describe backwards whiskers. The Cesàro equation describes how curvature changes along the whisker length. One disadvantage is that the coefficients A and B are very different across whiskers even within a single species.

Although it is true that any curve described by the Cesàro equation can be transformed to fit on the universal Euler spiral, it is not immediately clear what insights can be obtained by that transformation. A more significant advantage of the Cesàro fit may be that its parameters could represent how the whisker grows (from base to tip). Given that both whisker taper and intrinsic curvature are formed by the hair keratinization process, and that whiskers grow from base to tip, it is possible that the A and B coefficients of the Cesàro fits represent individual whisker growth. A promising hypothesis for future work is that whisker curvature and taper trade off with each other to normalize bending mechanics.

